# Growth-law offset and efficient protein synthesis in fast-growing bacteria

**DOI:** 10.64898/2026.11.17.767273

**Authors:** Hugo Dourado, Xiao-Pan Hu, Diana Széliová, Matteo Mori, Terence Hwa, Martin J. Lercher

## Abstract

Growth laws refer to simple relationships between various cellular components and the growth rate for exponentially growing bacteria subjected to various environmental and genetic perturbations. The best studied among these is the ribosomal growth law, which refers to a relationship between the ribosome content and the growth rate under different nutrient conditions. This relationship is affine, which means a linear dependence on the growth rate plus a constant offset. This offset, referred to here as the “growth-law offset” and which is substantial in size, has been rationalized in terms of ribosome sequestration and protein degradation effects in the slow growth regime. These effects however cannot account for the sizable offset observed at medium and fast growth, where the efficiency of protein synthesis is predominant. Here, we use a simple effective model of translation to show that efficient protein synthesis in the medium to fast-growth regime does in fact require a substantial growth-law offset, on the order of that observed in Escherichia coli. Our key observation is that efficient protein synthesis requires a nonlinear relationship between non-sequestered ribosomes and growth rate, but in the medium to fast-growth regime this relationship is well approximated by the affine relation in the growth law, with the offset being necessary for the efficient balance between ribosomal and non-ribosomal proteins directly involved in protein synthesis.

---

The ribosome content in bacterial cells growing under different levels of nutrient limitation shows an approximate affine dependence on the growth rate, a relationship known as the ribosomal growth law (1). An affine relation consists of a linear term plus a constant offset, in this case, expressed as

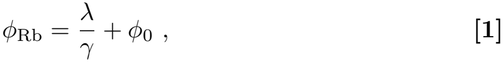

where *ϕ*_Rb_ is the proteome mass fraction of ribosomal proteins (hereafter also referred to as the Rb fraction), *λ* is the growth rate, and *γ* and *ϕ*_0_ are positive constants. The constant *γ* has been associated with an approximately constant elongation rate in bacteria (1), while the offset *ϕ*_0_ is about 5% of the proteome. At slow growth, the offset has been attributed to “inactive” ribosomes (2), which can arise mechanistically from ribosome sequestration (3) or as additional ribosome necessary to compensate for protein degradation (4); the benefit of withholding ribosomes as reserve at slow growth has been rationalized in the context of rapid transition to fast growth when nutrient condition improves (5). These effects cannot, however, explain the offset extrapolated from data at higher growth rates (e.g. higher than 0.5 h ^−1^), where sequestered ribosomes represent only 1% of the Rb proteins (2), protein degradation is approximately 20- to 100-fold slower than dilution by growth (4), and ribosome reserves for future environments (5) would tend to decrease in at least some bacteria well adapted to a particular steady growth condition. The offset *ϕ*_0_ thus represents an apparent inefficiency at higher growth rates.

Here we combine a simple effective model of translation based on Ref. (6) and our previous considerations on the efficient balance of enzymes and substrates (7), to show that the growth-law offset is in fact compatible with an efficient protein synthesis at the maximal growth rate.

## Results

### Efficient protein synthesis in a simple effective model of translation

We revisit the coarse-grained proteome partitioning model introduced by Klumpp et al. (6), but simplify it by accounting only for the proteome mass fraction *ϕ*_R_ directly involved in translation

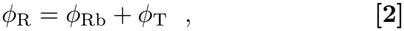

where *ϕ*_T_ denotes here the proteome fraction of tRNA-affiliated proteins devoted to ternary-complex supply, here comprising EF-Tu and the aminoacyl-tRNA synthetases. Following Ref. (6), we also assume that protein synthesis i) is catalyzed by Rb proteins, ii) has an elongation rate *ϵ*(*ϕ*_T_) that is a function of *ϕ*_T_ according to the Michaelis-Menten rate law, and iii) conserves mass without protein degradation. Differently from Ref. (6), however, we assume no fixed amount of sequestered ribosomes, and here *ϵ* is to be understood as an apparent elongation rate relative to all Rb proteins. These assumptions result in a direct expression for the growth rate *λ* as a function of *ϕ*_Rb_ and *ϕ*_T_ (see SI Appendix, extended methods)

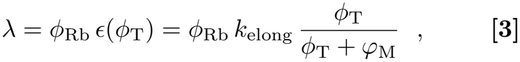

where the constant *k*_elong_ is the maximal peptide elongation rate per ribosomal protein, and *φ*_M_ is a type of Michaelis constant (6). Notably, this simple model fails at growth rates below about 0.5 h ^−1^, where the fraction of sequestered ribosomes becomes significant and the elongation rate per active ribosome does not go to zero (2, 3). Our analysis therefore focuses mainly on the regime of medium to fast growth at *λ >* 0.5 h^−1^, where the Michaelis-Menten type of relationship in Eq. 3 still holds reasonably well (2, 6).

We now combine this simple effective model with our previous considerations about the efficient balance of enzymes and substrates (7): we look for the most efficient combination of *ϕ*_Rb_, *ϕ*_T_ at any given proteome investment *ϕ*_R_ in Eq. 2 that results in the maximal growth rate *λ* in Eq. 3 (or equivalently, the minimal investment *ϕ*_R_ for any given *λ*, see SI Appendix, extended methods), which provides the most efficient *ϕ*_Rb_, *ϕ*_T_ as functions of *λ*

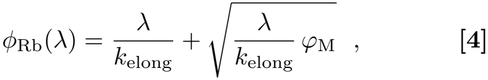

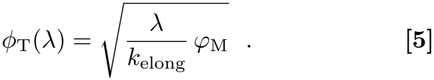

We observe that the efficient Rb fraction *ϕ*_Rb_ in Eq. 4 has two limiting regimes with respect to *λ*: one at low *λ* where the square-root term dominates the expression, and one at higher *λ* where the term *λ/k*_elong_ dominates the expression and *ϕ*_Rb_ is approximately linear in *λ*, with a smaller contribution from the square-root term. This means the affine relation in the growth law Eq. 1 matches qualitatively the efficient *ϕ*_Rb_ at higher growth rates, motivating the following quantitative analysis.

### Model parameterization and growth law from linear expansion

Fig. 1A shows the predictions of Eq. 4 for *E. coli* (red line) using the parameters *k*_elong_ = 12.7 h^−1^1 (corresponding to the estimated maximal elongation rate of 22 amino acids per second per ribosome (2), see Methods), and *φ*_M_ = 0.050 obtained by matching Eq. 5 to the independent T fraction regression extrapolated to maximal growth in Fig. 1B (see SI Appendix, extended methods). In Fig. 1A, we observe that at higher growth rates *λ >* 0.5 h^−1^1 the model approximates well the growth law resulting from a linear regression (dashed line) of the available data.

**Fig. 1.**
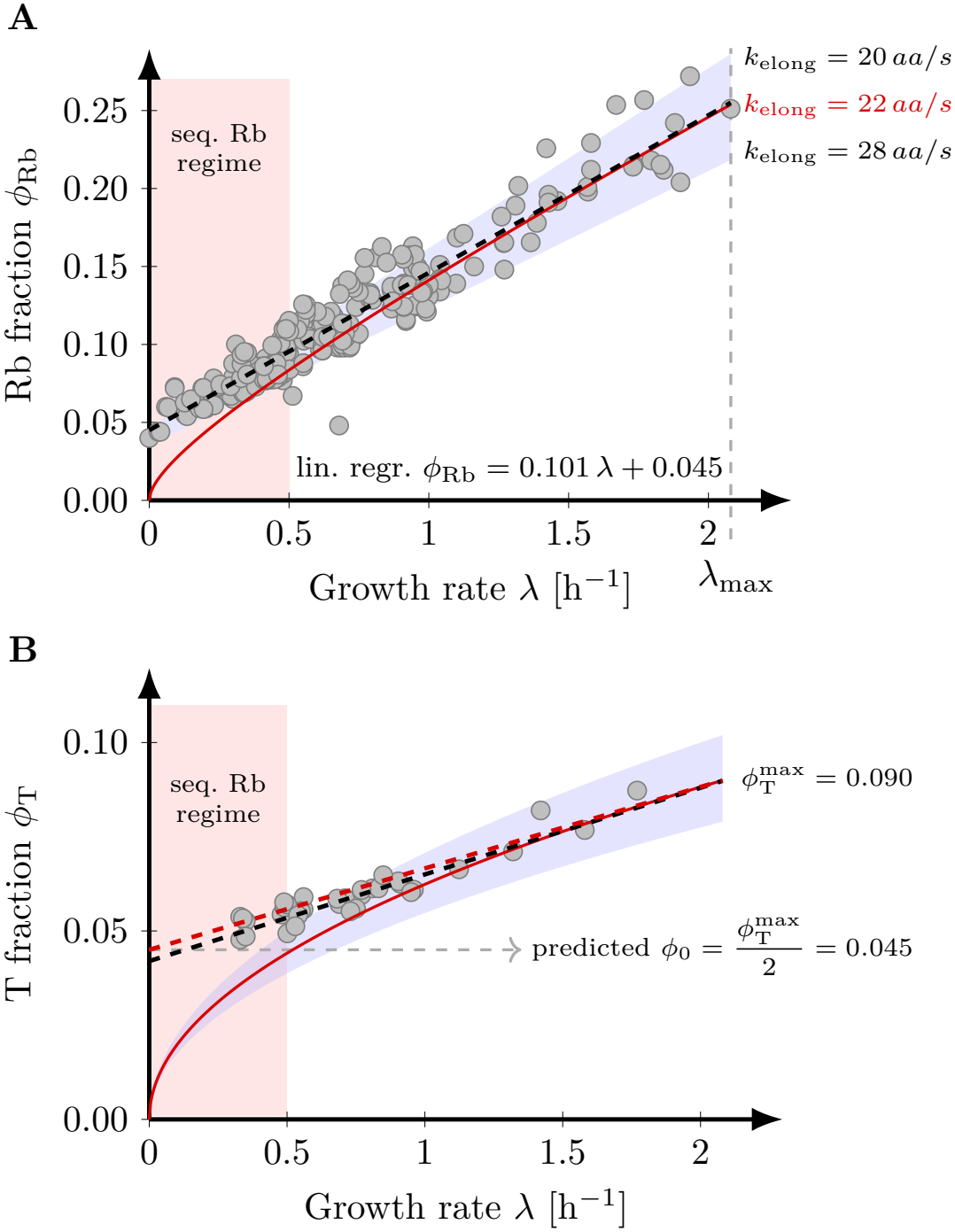
Efficient protein synthesis model (Eqs. 4,5) compared with *E. coli* data across nutrient qualities. Red regions indicate the regime outside the model’s scope, where ribosome sequestration (2, 3) and protein degradation (4) become relevant. Black dashed lines indicate linear regressions of the data. A) Eq. 4 (red line), with *k*_elong_ = 22 aa s^−1^ and *φ*_M_ = 0.050, approximates the linear regression at maximal growth rate. The parameter *φ*_M_ matches Eq. 5 to the independent T data regression in B at *λ*_max_ = 2.08 h^−1^. Blue shading spans linear expansions at *λ*_max_ for the indicated *k*_elong_ values. B) Eq. 5 (red line), with *φ*_M_*/k*_elong_ = 0.0039 h, is consistent with the T data (8, 9) (gray) in the fast growth regime. The red dashed line indicates the linear expansion of Eq. 5 at *λ*_max_; blue shading spans predictions for *φ*_M_*/k*_elong_ = 0.003–0.005 h. The predicted offset *ϕ*_0_ = 0.045 equals half the estimated 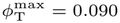 at *λ*_max_.

We formalize this quasi-linear regime of Eq. 4 at higher growth rates by taking its linear expansion at the maximal growth rate *λ*_max_ = 2.08 h^−1^1 observed in *E. coli* (10), resulting in an affine relation as the growth law in Eq. 1, but with the offset and slope determined respectively as (see SI Appendix, extended methods)

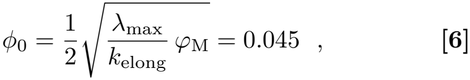

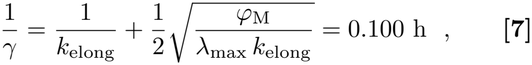

which are in agreement with the offset 0.045 *±* 0.002 and slope 0.101 *±* 0.002 h (estimate *±* SE) obtained from the linear regression of the data in Fig. 1A. This good quantitative agreement is also robust to some variation in the model parameters; e.g. a change of 30% in an individual parameter used to predict the offset in Eq. 6 (including the maximal growth rate *λ*_max_) would cause less than 20% change in the offset prediction due to the square-root dependency. The predicted slope in Eq. 7 has an additional dependency on the *k*_elong_ parameter, but some variation here still provides reasonable predictions; see the blue region in Fig. 1A for *k*_elong_ from 20 to 28 aa/s, as a reference. We note that the parameter *φ*_M_ = 0.050 used to predict the growth law in Fig. 1A was fitted to the independent T fraction data in Fig. 1B. This value lies in a physically reasonable range, between the estimated lower bound of *φ*_M_ *≈* 0.02 imposed by diffusion-limited binding of ternary complexes (6) and the upper bound *φ*_M_ *≈* 0.075 set by molecular crowding at *λ*_max_ *≈* 2.08 h^−1^1 according to the model of Klumpp et al. (6).

The model also captures two non-trivial properties observed in the data. First, by combining Eqs. 5, 6 we find that the model predicts the offset *ϕ*_0_ must be half of the T fraction at maximal growth rate 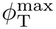

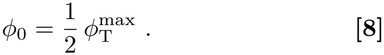

Eq. 8 is consistent with the value 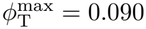 obtained by extrapolating the linear regression of the T fraction data in Fig. 1B to *λ*_max_ = 2.08 h^−1^1, and the Rb offset *ϕ*_0_ = 0.045. The second non-trivial property comes from the linear expansion of Eq. 5 around the maximum growth rate, for which we find an offset also determined by Eq. 6 (see SI Appendix, extended methods). This means the model predicts that both Rb and T fractions should have similar offsets when extrapolating the linear regression of their fast-growth data to zero growth, which is consistent with the offset 0.042 *±* 0.001 resulting from the linear regression of the T fraction data in Fig. 1B.

Finally, we note that quantitative precision should neither be expected from nor overinterpreted in such a simple effective model, whose contribution is mainly conceptual: an efficient balance between Rb and T proteins near maximal growth in *E. coli* requires a significant Rb offset, on the order of 5% of the proteome as observed in empirical growth laws.

## Discussion

Here, we show that the growth-law offset is compatible with an efficient investment in ribosomal proteins (Rb-protein) at higher growth rates, and that its value under efficient allocation equals half of the tRNA-affiliated proteins (T-protein) at maximal growth. This result brings together our previous findings that efficient enzyme investment follows a linear-plus-square-root dependence on flux (7), that a minimal ribosome-substrate model predicts the nonlinear dependence of active ribosomes on growth rate in *E. coli* and yeast (11), and that a detailed kinetic model reproduces the measured composition of the *E. coli* translation machinery under the assumption of efficient protein synthesis (12). Here, we isolate the minimal quantitative principle connecting these results to the growth-law offset by expanding a two-component model of Rb and T proteins around the maximal growth rate. This result differs from previous studies that associate the offset with a fixed pool of sequestered ribosomes also at high growth rates (1) and that derive similar nonlinear relations based on models with more proteome components (6, 13).

Our results thus clarify the quantitative relationship between the growth-law offset and the efficiency of protein synthesis at fast growth, complementing previous studies explaining how growth laws arise through regulatory mechanisms in *E. coli* (3) and how ribosome sequestration account for the slow-growth offset (2). Our minimal model does not, however, explain why the offset extrapolated from fast growth agrees so well with this slow-growth offset or, more generally, why the growth law remains approximately affine across all growth regimes. As a possibility for future work, we speculate that both observations may still be compatible with efficient translation in an extended model incorporating the constraint imposed by the ppGpp-mediated regulatory mechanism (3).

## Materials and Methods

### Unit conversion for *k*_elong_

Following Ref. (11), we convert *k*_elong_ from aa s^−1^ to protein mass synthesized per ribosomal protein mass per hour using an average amino acid molar mass of 132 g mol^−1^, a ribosome molar mass of 2,306,967 g mol^−1^, and a ribosomal protein mass fraction 0.358. The conversion factor is 132 *×* 3600*/*(2,306,967 *×* 0.358) = 0.575 h ^−1^.

### Data collection

For Fig. 1A, we used the compilation of Chure and Cremer (13), supplemented with an LB-growth data point from Ref. (10) and data from Refs. (8, 9, 14). We excluded the data of Si et al. (15) due to its substantially higher variance, but its inclusion results in an offset 0.049 *±* 0.002, still at the order of 5% of the proteome. The data in Fig. 1B were obtained from Refs. (8, 9).

### Data, Materials, and Software Availability

All processed data and analysis code are available at https://github.com/diana-sz/offset_data.

## ACKNOWLEDGMENTS

This work was funded through grants to M.J.L. by the Deutsche Forschungsgemeinschaft (DFG, German Research Foundation: CRC 1310), and to D.S. by the Austrian Science Fund (FWF) 10.55776/J4858.

