## Supporting Information for "Growth-law offset and efficient protein synthesis in fast-growing bacteria"

### Supporting Information Text

### Extended methods

**Derivation of the growth rate $\lambda$ as a function of $\phi_{\mathrm{Rb}}$ and $\phi_{\mathrm{T}}$.** Our simple effective model of translation assumes that the translation reaction at steady-state growth (also known as balanced growth (1)) has a flux $v$ in $\mathrm{g\,L^{-1}\,h^{-1}}$ which is catalyzed by Rb proteins with corresponding proteome mass fraction $\phi_{\mathrm{Rb}}$ of the total proteome mass concentration $c_{\mathrm{P}}$ in $\mathrm{g\,L^{-1}}$, and with elongation rate per ribosomal protein mass $\epsilon$ in $\mathrm{h^{-1}}$

$$v = \phi_{\mathrm{Rb}}\, c_{\mathrm{P}}\, \epsilon \quad . \tag{1}$$

We also assume the elongation rate follows the Michaelis-Menten rate law

$$\epsilon(\phi_{\mathrm{T}}) = k_{\mathrm{elong}} \frac{\phi_{\mathrm{T}}}{\phi_{\mathrm{T}} + \varphi_{\mathrm{M}}} \quad , \tag{2}$$

with $k_{\mathrm{elong}}$ being the maximal elongation rate of peptide mass per ribosomal protein mass in $\mathrm{h^{-1}}$ unit, and $\varphi_{\mathrm{M}}$ is a dimensionless type of Michaelis constant as previously defined in Ref. (2).

Finally, we assume the mass conservation of protein synthesis flux without protein degradation, which means all protein synthesis flux goes into the dilution of proteins by growth at rate $\lambda\, c_{\mathrm{P}}$

$$v = \lambda\, c_{\mathrm{P}} \quad . \tag{3}$$

Combining Eqs. 1,2,3 and canceling the $c_{\mathrm{P}}$ term, we get Eq. 3 in the main text.

**Derivation of the most efficient $\phi_{\mathrm{Rb}}$, $\phi_{\mathrm{T}}$ as functions of $\lambda$.** Here we define the most efficient proteome mass fractions $\phi_{\mathrm{Rb}}$, $\phi_{\mathrm{T}}$ as the solution to the problem of getting the highest growth rate $\lambda$ in the main text's Eq. 3 with the constraint given by the sum $\phi_{\mathrm{R}} = \phi_{\mathrm{Rb}} + \phi_{\mathrm{T}}$ in main text's Eq. 2. We see next that this solution does not require any particular value of $\phi_{\mathrm{R}}$ given as input, and that it is also the solution that minimizes the sum $\phi_{\mathrm{R}} = \phi_{\mathrm{Rb}} + \phi_{\mathrm{T}}$ under the constraint of Eq. 3 in the main text. We first combine both Eqs. 2,3 in the main text and solve it for $\lambda$ as a function of $\phi_{\mathrm{T}}$ only

$$\lambda(\phi_{\mathrm{T}}) = k_{\mathrm{elong}} \frac{(\phi_{\mathrm{R}} - \phi_{\mathrm{T}})\phi_{\mathrm{T}}}{\phi_{\mathrm{T}} + \varphi_{\mathrm{M}}}. \tag{4}$$

For $\phi_{\mathrm{R}} > 0$, $k_{\mathrm{elong}} > 0$, and $\varphi_{\mathrm{M}} > 0$, this expression is positive for $0 < \phi_{\mathrm{T}} < \phi_{\mathrm{R}}$ and vanishes at both endpoints. Its derivative is

$$\frac{d\lambda}{d\phi_{\mathrm{T}}} = k_{\mathrm{elong}} \frac{\phi_{\mathrm{R}}\varphi_{\mathrm{M}} - 2\varphi_{\mathrm{M}}\phi_{\mathrm{T}} - \phi_{\mathrm{T}}^2}{(\phi_{\mathrm{T}} + \varphi_{\mathrm{M}})^2}. \tag{5}$$

The numerator decreases strictly with $\phi_{\mathrm{T}}$ over the allowed interval, changing sign from positive to negative. Its unique zero therefore gives the global maximum:

$$\phi_{\mathrm{R}}\varphi_{\mathrm{M}} = \phi_{\mathrm{T}}^2 + 2\varphi_{\mathrm{M}}\, \phi_{\mathrm{T}}. \tag{6}$$

Using $\phi_{\mathrm{R}} = \phi_{\mathrm{Rb}} + \phi_{\mathrm{T}}$, this condition becomes

$$\phi_{\mathrm{Rb}} = \phi_{\mathrm{T}} + \frac{\phi_{\mathrm{T}}^2}{\varphi_{\mathrm{M}}} \ . \tag{7}$$

Substituting this relation into Eq. 3 of the main text gives

$$\lambda = k_{\mathrm{elong}} \frac{\phi_{\mathrm{T}}^2}{\varphi_{\mathrm{M}}} \ . \tag{8}$$

Now solving for $\phi_{\mathrm{T}}$, we get the Eq. 5 in the main text, representing the efficient fraction $\phi_{\mathrm{T}}(\lambda)$ in terms of the growth rate $\lambda$ without having to specify $\phi_{\mathrm{R}}$. Now substituting Eq. 5 of the main text into Eq. 7 above, we get Eq. 4 in the main text.

Equivalently, the elongation rate defined in Eq. 2 in terms of the efficient $\phi_{\mathrm{T}}$ defined by Eq. 5 in the main text becomes

$$\epsilon(\lambda) = \frac{k_{\mathrm{elong}}}{1 + \sqrt{\dfrac{k_{\mathrm{elong}}\, \varphi_{\mathrm{M}}}{\lambda}}} \ . \tag{9}$$

So the inverse slope $\gamma$ at the maximal growth rate is the harmonic mean between $k_{\mathrm{elong}}$ and the maximal elongation rate $\epsilon_{\max} = \epsilon(\lambda_{\max})$

$$\gamma = \frac{2\, k_{\mathrm{elong}}\, \epsilon_{\max}}{k_{\mathrm{elong}} + \epsilon_{\max}} \ . \tag{10}$$

To establish the equivalent minimization formulation, we now fix $\lambda > 0$ and solve Eq. 3 of the main text for $\phi_{\mathrm{Rb}}$. The total investment is then

$$\phi_{\mathrm{R}}(\phi_{\mathrm{T}}) = \phi_{\mathrm{T}} + \frac{\lambda}{k_{\mathrm{elong}}} + \frac{\lambda\,\varphi_{\mathrm{M}}}{k_{\mathrm{elong}}\,\phi_{\mathrm{T}}} \ . \quad [11]$$

Its first and second derivatives are

$$\frac{d\phi_{\mathrm{R}}}{d\phi_{\mathrm{T}}} = 1 - \frac{\lambda\,\varphi_{\mathrm{M}}}{k_{\mathrm{elong}}\,\phi_{\mathrm{T}}^2}, \quad [12]$$

$$\frac{d^2\phi_{\mathrm{R}}}{d\phi_{\mathrm{T}}^2} = \frac{2\lambda\,\varphi_{\mathrm{M}}}{k_{\mathrm{elong}}\,\phi_{\mathrm{T}}^3} > 0. \quad [13]$$

The unique minimum therefore occurs at $\phi_{\mathrm{T}} = \sqrt{\lambda\varphi_{\mathrm{M}}/k_{\mathrm{elong}}}$, recovering the same efficient fractions. Maximizing growth at fixed investment and minimizing investment at fixed growth thus identify the same relationship between $\phi_{\mathrm{Rb}}$, $\phi_{\mathrm{T}}$, and $\lambda$.

**Fitting the parameter $\varphi_{\mathrm{M}}$ to the T fraction data at the maximal growth rate.** Here we estimate the ratio $\varphi_{\mathrm{M}}/k_{\mathrm{elong}}$ in Eq. 5 of the main text by requiring the predicted T fraction at maximal growth to match the linear regression of the T fraction data evaluated at the same growth rate.

We first calculate the linear regression of the data points shown in Fig. 1B of the main text by ordinary least squares using the R function `lm` (3)

$$\phi_{\mathrm{T}}^{\mathrm{regression}}(\lambda) = (0.023\ \mathrm{h})\lambda + 0.042 \ . \quad [14]$$

Extrapolating this relation to $\lambda_{\max} = 2.08\,\mathrm{h}^{-1}$ gives

$$\widehat{\phi_{\mathrm{T}}^{\max}} \simeq 0.090. \quad [15]$$

We then require Eq. 5 in the main text to reproduce this estimated fraction

$$\sqrt{\lambda_{\max}\frac{\varphi_{\mathrm{M}}}{k_{\mathrm{elong}}}} = \widehat{\phi_{\mathrm{T}}^{\max}}. \quad [16]$$

Solving for the parameter ratio gives

$$\frac{\varphi_{\mathrm{M}}}{k_{\mathrm{elong}}} = \frac{\left(\widehat{\phi_{\mathrm{T}}^{\max}}\right)^2}{\lambda_{\max}} \simeq 0.0039\,\mathrm{h}. \quad [17]$$

Combining this ratio with $k_{\mathrm{elong}} = 12.7\,\mathrm{h}^{-1}$ gives $\varphi_{\mathrm{M}} \simeq 0.050$. With these values $k_{\mathrm{elong}} = 12.7\,\mathrm{h}^{-1}$ and $\varphi_{\mathrm{M}} = 0.050$, we predict the following offset and slope from Eqs. 6,7 in the main text

$$\phi_0 = \frac{1}{2}\sqrt{\frac{\lambda_{\max}\,\varphi_{\mathrm{M}}}{k_{\mathrm{elong}}}} \simeq 0.045 \ , \quad [18]$$

$$\frac{1}{\gamma} = \frac{1}{k_{\mathrm{elong}}} + \frac{1}{2}\sqrt{\frac{\varphi_{\mathrm{M}}}{\lambda_{\max}\,k_{\mathrm{elong}}}} \simeq 0.100\,\mathrm{h} \ . \quad [19]$$

These values match the offset $0.045 \pm 0.002$ and slope $0.101 \pm 0.002$ h (estimate ± SE) from a linear regression of the Rb fraction data points shown in Fig. 1A of the main text, by ordinary least squares using the R function `lm` (3).

**Linear expansion of Eq. 4 in the main text at the maximal growth rate.** Here we take the linear expansion of Eq. 4 in the main text around the maximal growth rate $\lambda_{\max}$. We first calculate its derivative with respect to $\lambda$

$$\frac{d\phi_{\mathrm{Rb}}}{d\lambda} = \frac{1}{k_{\mathrm{elong}}} + \frac{1}{2}\sqrt{\frac{\varphi_{\mathrm{M}}}{\lambda k_{\mathrm{elong}}}}. \quad [20]$$

The linear expansion is then

$$\phi_{\mathrm{Rb}}(\lambda) \simeq \phi_{\mathrm{Rb}}(\lambda_{\max}) + \left.\frac{d\phi_{\mathrm{Rb}}}{d\lambda}\right|_{\lambda_{\max}} (\lambda - \lambda_{\max}). \quad [21]$$

Substituting Eq. 4 in the main text and the derivative above, we get

$$\phi_{\mathrm{Rb}}(\lambda) \simeq \frac{\lambda_{\max}}{k_{\mathrm{elong}}} + \sqrt{\frac{\lambda_{\max}\,\varphi_{\mathrm{M}}}{k_{\mathrm{elong}}}} + \left(\frac{1}{k_{\mathrm{elong}}} + \frac{1}{2}\sqrt{\frac{\varphi_{\mathrm{M}}}{\lambda_{\max}\,k_{\mathrm{elong}}}}\right)(\lambda - \lambda_{\max}). \quad [22]$$

Now collecting the terms proportional to $\lambda$ and the constant terms separately, we get

$$\phi_{\mathrm{Rb}}(\lambda) \simeq \left(\frac{1}{k_{\mathrm{elong}}} + \frac{1}{2}\sqrt{\frac{\varphi_{\mathrm{M}}}{\lambda_{\max}\,k_{\mathrm{elong}}}}\right)\lambda + \frac{1}{2}\sqrt{\frac{\lambda_{\max}\,\varphi_{\mathrm{M}}}{k_{\mathrm{elong}}}}. \quad [23]$$

This is the same affine relationship as the growth law $\phi_{\mathrm{Rb}} = \lambda/\gamma + \phi_0$ in the main text's Eq. 1, with offset and slope given by Eqs. 6,7 in the main text.

**Linear expansion of Eq. 5 in the main text at the maximal growth rate.** Here we take the linear expansion of Eq. 5 in the main text around the maximal growth rate $\lambda_{\max}$. We first calculate its derivative with respect to $\lambda$

$$\frac{d\phi_{\mathrm{T}}}{d\lambda} = \frac{1}{2}\sqrt{\frac{\varphi_{\mathrm{M}}}{\lambda\, k_{\mathrm{elong}}}}. \quad [24]$$

The linear expansion is then

$$\phi_{\mathrm{T}}(\lambda) \simeq \phi_{\mathrm{T}}(\lambda_{\max}) + \left.\frac{d\phi_{\mathrm{T}}}{d\lambda}\right|_{\lambda_{\max}} (\lambda - \lambda_{\max}). \quad [25]$$

Substituting Eq. 5 in the main text and the derivative above, we get

$$\phi_{\mathrm{T}}(\lambda) \simeq \sqrt{\frac{\lambda_{\max}\,\varphi_{\mathrm{M}}}{k_{\mathrm{elong}}}} + \frac{1}{2}\sqrt{\frac{\varphi_{\mathrm{M}}}{\lambda_{\max}\, k_{\mathrm{elong}}}}(\lambda - \lambda_{\max}). \quad [26]$$

Now collecting the terms proportional to $\lambda$ and the constant terms separately, we get

$$\phi_{\mathrm{T}}(\lambda) \simeq \frac{1}{2}\sqrt{\frac{\varphi_{\mathrm{M}}}{\lambda_{\max}\, k_{\mathrm{elong}}}}\lambda + \frac{1}{2}\sqrt{\frac{\lambda_{\max}\,\varphi_{\mathrm{M}}}{k_{\mathrm{elong}}}}. \quad [27]$$

We see that the offset is the same as $\phi_0$ in the linear expansion of $\phi_{\mathrm{Rb}}$ derived above. Using Eq. 5 in the main text at $\lambda_{\max}$, we also get

$$\phi_0 = \frac{1}{2}\sqrt{\frac{\lambda_{\max}\,\varphi_{\mathrm{M}}}{k_{\mathrm{elong}}}} = \frac{1}{2}\phi_{\mathrm{T}}^{\max}, \quad [28]$$

which is Eq. 8 in the main text. Thus, the linear expansions of both fractions have the same offset, equal to half of the efficient T fraction at maximal growth.

**Analysis of growth law and T fraction data from literature.** For Fig. 1A, showing the $\phi_{\mathrm{Rb}}$ fraction (ribosomal proteins), we used the compilation of Chure and Cremer (4). We excluded the data of Si et al. (5) because of their substantially higher variance. We added an LB-growth data point from Ref. (6). For the datasets in (7, 8), we extracted the ribosomal protein fraction from proteomics data by summing over the following ribosomal genes:

- rplA, rplB, rplC, rplD, rplE, rplF, rplI, rplJ, rplK, rplL, rplM, rplN, rplO, rplP, rplQ, rplR, rplS, rplT, rplU, rplV, rplW, rplX, rplY, rpmA, rpmB, rpmC, rpmD, rpmE, rpmF, rpmG, rpmH, rpmI, rpmJ, rpsA, rpsB, rpsC, rpsD, rpsE, rpsF, rpsG, rpsH, rpsI, rpsJ, rpsK, rpsL, rpsM, rpsN, rpsO, rpsP, rpsQ, rpsR, rpsS, rpsT, rpsU, sra,

taken from Supplementary Table 3 of Wu et al. 2023 (8). Protein abundances in each dataset were first normalized to sum to 1 across the proteome, then summed over this gene set to obtain $\phi_{\mathrm{Rb}}$. We used conditions from carbon-limited chemostat and minimal media from (7) and the steady-state samples in (8). From (9), we multiplied the reported RNA/protein ratios by 0.5 to obtain the ribosomal protein fraction.

Fig. 1B, showing the $\phi_{\mathrm{T}}$ fraction, is defined here as the combined proteome mass fraction of EF-Tu and aminoacyl-tRNA synthetases, comprising the following genes:

- Elongation factor (EF-Tu): tufA, tufB;
- Aminoacyl-tRNA synthetases: alaS, argS, asnS, aspS, cysS, glnS, gltX, glyQ, glyS, hisS, ileS, leuS, lysS, lysU, metG, pheS, pheT, proS, serS, thrS, trpS, tyrS, valS.

The aminoacyl-tRNA synthetase gene set was taken from Supplementary Table 3 of Wu et al. 2023 (8). For each gene, the corresponding protein abundance was obtained from the proteomics datasets (7, 8), normalized as described above and for the same conditions.
